# meTCRs - Learning a metric for T-cell receptors

**DOI:** 10.1101/2022.10.24.513533

**Authors:** Felix Drost, Lennard Schiefelbein, Benjamin Schubert

**Affiliations:** Computational Health Center, Helmholtz Munich Neuherberg, Germany

## Abstract

T cell receptors (TCRs) bind to pathogen- or self-derived epitopes to elicit a T cell response as part of the adaptive immune system. Determining the specificity of TCRs provides context for immunological studies and can be used to identify candidates for novel immunotherapies. To avoid costly experiments, large-scale TCR-epitope databases are queried for similar sequences via various distance functions. Here, we developed the deep-learning based distance *meTCRs*. Contrary to most previous approaches, the method avoids computational expansive pairwise string operations by comparing TCRs in a numeric embedding. In contrast to models which are trained specificity-agnostic, we directly utilize epitope information by applying deep metric learning to guide the training. Summarizing, we present *meTCRs* as a scalable alternative to embed TCR repertoires for clustering, visualisation, and querying against the ever-increasing amount TCR-epitope pairs in publicly available databases.

## 1 Introduction

T cells recognize epitopes, i.e., peptide sequences derived from antigens, via the highly variable T cell receptor (TCR). Due to the immense diversity, the adaptive immune system is able to fight a wide variety of diseases including infections and tumors: At least 10^7^ different TCR sequences exist in an individual human being, while population wide estimation exceed 10^20^ TCRs [1]. Through recent advances in single-cell technology, T cell specificity can now be determined in large TCR repertoires for predefined epitopes [2]. Early studies have shown, that TCRs with similar sequences are likely bind to the same epitope [3, 4]. Following, various approaches were developed to cluster TCR sequences based on their specificity, or to perform database queries by calculating pairwise sequence similarity. Most approaches are either based on string-based comparison [5, 6, 7] or numeric embeddings [8, 9]. Typically, string-base approaches make use of heuristic similarity measurements, which often rely of computationally costly pairwise string operations. While this is partially avoided for numeric representations, they usually rely on unsupervised approaches such as biophysical-informed embeddings or Variational Autoencoders, which are not specifically trained to preserve epitope specificity. To circumvent these shortcomings, we propose *meTCRs* - a numeric TCR embedding, which incorporates specificity annotation during training by applying supervised deep metric learning [10]. We envision *meTCRs* to become a valuable tool to derive meaningful representations of TCRs, which seamlessly scale to downstream tasks such as clustering on large-scale single-cell datasets or querying vast databases to assign specificity. Furthermore, the method can continuously improve by incorporating novel developments in the field of deep metric learning and by the ever-increasing amount of publicly available training data.

## 2 Method

### Data pre-processing

Paired epitope and TCR sequencing data were obtained from three publicly available databases IEDB [11], VDJdb [12], and McPAS-TCR [13] for training and validation. More specifically, we extracted the hypervariable complementarity-determining region 3 of the beta-chain (CDR3-*β*) of the TCR as it lays in close proximity to the epitope and is assumed to be the driving factor in antigen recognition [14] and is therefore most prevalent in those databases. To harmonize the datasets, leading cysteine and ending phenylalanine or tryptophan were removed from the CDR3-*β* sequences. The amino acid sequences were end-padded to the maximal sequence length (*l* = 36) in the dataset and one-hot encoded. The epitope annotations were solely used as categorical labels.

### Architecture

Let 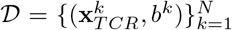 be a dataset consisting of the pre-processed CDR3-*β* sequence 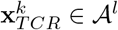 and the categorical binding label *b*^*k*^ ∈ B for each database entry *k*, where A represents the set of possible amino acids and padding character, and B corresponds to the set of all binding epitopes. *meTCRs* consists of an embedding model *E*, which takes the 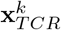 as an input. As embedding model, we implemented multilayer perceptrons (MLPs), convolutional neural networks (CNNs), long short-term memory networks (LSTMs) [15], and transformers [16]. For the first three architectures the one-hot encoded CDR3-*β* sequence is directly used as input. For the transformer encoder, an additional position-wise encoding was added to a learnt embedding layer. Eventually, a final linear layer is applied to the output of the last network layer. Thereby, the embedding network projects the input data into the representation 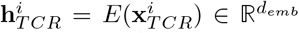 with the embedding dimension *d*_*emb*_ on which all downstream tasks are performed.

### Loss function

Most deep metric learning losses are calculated on batches of both positive and negative pairs [10]. Typically, selecting informative tuples during training is a challenging design choice. In the case of *meTCRs*, a pair of TCRs is a positive pair if the receptors of database entry *A* and *B* bind on a common epitope (*b*^*A*^ = *b*^*B*^) and negative if this is not the case (*b*^*A*^≠*b*^*B*^). However, common TCR databases only contain experimentally validated epitope bindings and miss negative pairs. Furthermore, TCRs might be cross-reactive, i.e., a TCR may recognize multiple epitopes. Therefore, randomly creating samples by pairing TCRs from different epitope classes can create false-negative pairs. The Barlow Twin loss

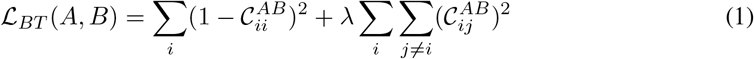

circumvents these challenges [17]. Here, the cross correlation matrix

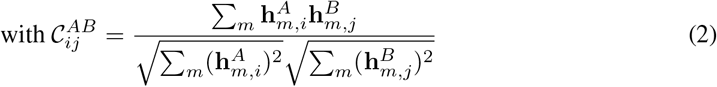

of batch-normalized inputs is calculated on positive pairs only. Overall, the Barlow Twin loss maximizes the similarity between paired representation while the embedding’s redundancy is minimized. Batch normalization ensured that no trivial representation is learned even though only positive pairs are used.

### Training Details

The models were trained on the processed data from IEDB and VDJdb excluding duplicates, which resulted in N=145,331 entries. Pairs of 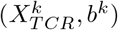 were randomly divided into training and validation set with a split of 80%-20%. Due to the nature of the Barlow Twin loss, the batches need to be composed of paired input sequences. To this end, epitope classes which contained only one TCR sequence within a set were removed. During training, hierarchical sampling is applied to build batches of size *N*. In the first step, 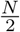 epitopes are sampled randomly and proportional to the epitope distribution within the training set to prevent over-representation of small classes. Following, two binding TCRs are sampled for all selected epitopes. The model was trained using the ADAM optimizer [18]. All hyperparameter of the embedding model, the optimizer, and the loss function were optimized via the Tree-Structured Parzen Estimator (TPE) algorithm [19] as implemented in the Optuna framework [20] to maximize the mean average precision (MAP) [10] at 10 on the validation set (see Results).

## 3 Results

We generated an independent set from the processed McPAS-TCR database by removing all pairs which already appear in the training or validation set from the IEDB and VDJdb and epitopes with only one reported TCR, which resulted in a test set 𝒯of N=7,733. Note, that prediction on this set are especially challenging due to a shift in epitopes between databases. Only two out of the five most abundant epitopes in the training set are present in the test set, and two out of the five largest classes in the test set are available in the training set with more than 50 TCRs. We embedded the test set via the different versions of the embedding network *E*, followed by calculating pairwise distances using the Euclidean norm. Additionally, pairwise distances are derived from *TCRmatch* as a string-based baseline method.

To measure the ability to predict specificity with *meTCRs*, we report classification based metrics. Here, the k-nearest neighbors 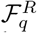 are determined for each element *q* of the test set based on the pairwise distances. Following, the metrics for F1-score (F1@R), precision (P@R), and recall (R@R) at *R* = 10 are reported. While R@R describes the average of samples for which a correct pair is contained within the *R* nearest neighbors, P@R is calculated as the average amount of matching samples within *R* nearest neighbors. F1@R is calculated as the harmonic mean between R@R and P@R. Further, we report the mean average precision at *R* ∈ {1, 10}

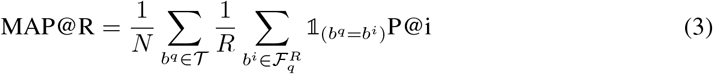

as proposed in [10] which incorporates the rank of the retrieval. To evaluate the consistency of *meTCRs’* embedding space, we cluster the embedded samples via the K-Means algorithm with K equal to the number of epitope classes in the test set and report the normalized mutual information (NMI). Finally, we report the AUC-ROC as a threshold based evaluation metric.

The choice of embedding network heavily influenced the models’ performance. The transformer-based model surpassed the remaining models by a large margin on all classification and clustering metrics (Table 1). However, *meTCRs* still falls short compared to the string-comparison based method *TCRmatch* by approximately one point across all metrics. In this evaluation, the predicted specificity by *meTCRs* and *TCRmatch* corresponds to the true specificity in approximately one third of the database queries. While this performance is not yet sufficient to accurately determine the specificity of TCRs, it provides researchers with a valuable initial guess for further experimental validation.

**Table 1:** Classification metrics of models trained with the IEDB-VDJdb data and TCRmatch on the McPAS-TCR dataset.

| Method | F1@10 | P@10 | R@10 | MAP@1 | MAP@10 | NMI |
| --- | --- | --- | --- | --- | --- | --- |
| MLP | 0.215 | 0.136 | 0.505 | 0.201 | 0.078 | 0.268 |
| CNN | 0.248 | 0.161 | 0.540 | 0.217 | 0.096 | 0.287 |
| LSTM | 0.209 | 0.133 | 0.486 | 0.146 | 0.072 | 0.271 |
| Transformer | 0.328 | 0.227 | 0.593 | 0.314 | 0.164 | **0.346** |
| TCRmatch | **0.336** | **0.232** | **0.609** | **0.330** | **0.170** | — |

Similar trends between the different embedding networks were also observed to a lesser degree for AUC-ROC based evaluation across the five most abundant epitopes (Table 2), except for the LSTM-based model, which surprisingly outperformed the transformer-model on the two most abundant epitopes and on the weighted average. Interestingly, *meTCRs* outperformed *TCRmatch* by large margins for most classes with at least one model version. Further, there is a large discrepancy in the performance for different epitopes. It needs to be further evaluated, whether the low similarity for these epitopes are inherent to the dataset and why different versions of the models fail to generalize to for individual classes.

**Table 2:** Support in the test (n-Test) and training set (n-Train) and AUC-ROC of models trained with the IEDB-VDJdb data and TCRmatch on the McPAS-TCR dataset for the five most abundant epitopes (long sequences were abbreviated).

| Epitope | n-Test | n-Train | MLP | CNN | LSTM | Transformer | TCRMatch |
| --- | --- | --- | --- | --- | --- | --- | --- |
| LPRRSGAAGA | 1,983 | 159 | 0.510 | 0.559 | **0.650** | 0.508 | 0.542 |
| CRVLCCYVL | 404 | 31 | 0.511 | 0.554 | **0.611** | 0.577 | 0.447 |
| WEDLFCDES... | 364 | 0 | 0.538 | 0.541 | 0.679 | **0.737** | 0.487 |
| GILGFVFTL | 358 | 3,502 | 0.435 | 0.435 | 0.472 | **0.528** | 0.485 |
| RFYKTLRAE... | 302 | 2 | 0.644 | 0.633 | 0.607 | **0.810** | 0.716 |
| Weighted AVG |  |  | 0.517 | 0.550 | **0.626** | 0.569 | 0.537 |

## 4 Discussion

Here, we introduced *meTCRs* for clustering and querying databases of TCRs as an alternative to classical string metrics or novel unsupervised approaches. Contrary to previous methods, *meTCRs* directly utilizes epitope binding information during training of a TCR distance by applying deep metric learning. The method can be easily scaled to large datasets or databases by pre-computing the numeric representation for each TCR, followed by the calculation of the computational cheap Euclidean pairwise distances on these embeddings.

In future work, the method can further be improved by adjusting for the imbalance in the databases by different sample schemes, loss weigthing, or further model adaptation. Exemplary, the IEDB, as largest source of TCR-epitope pairs, contains a total of 939 different classes. However, 49% of the TCRs stem from one of eight epitopes. Second, the performance can further be improved by collecting the ever-increasing amount of publicly available binding data. In previous studies, additional information on the TCR such as the *α*-chain, or V(D)J-gene annotation benefited TCR-epitope prediction and clustering [3]. However, full TCR-sequence data is sparse in public databases. Therefore, the model architecture could be adjusted for the missing value problem to embed TCRs with various degrees of information. Lastly, the presented evaluation highlights, that *meTCRs* generalizes well to unseen epitopes. However, an in-depth validation of *meTCRs* is yet to be performed for a wider variety of diseases and use cases. One major application of the method will be queries to databases, where most of the epitopes were already observed by the model during training. Therefore, *meTCRs* needs to be thoroughly compared to common methods like *TCRmatch* [6], *TCRdist* [5], and *DeepTCR* [8] for performance and computational cost in this use-case.

Upon these improvements, we envision *meTCRs* to become a fast and scalable solution for providing a specificity-informed representation of TCR repertoires that enables downstream tasks such as visualisation and clustering. Especially the ability to conduct computationally efficient queries to large TCR-epitope databases, provides a compelling alternative to determine specificity of TCR sequences from low-throughput experiments and large-scale single-cell studies without the need for exhaustive wet-lab experiments.

## Data and code availability

The corresponding data are publicly available and were downloaded from IEDB (https://www.iedb.org/, accessed March, 6^th^, 2022), VDJdb (https://vdjdb.cdr3.net/, accessed February, 25^th^, 2022), and McPAS-TCR (http://friedmanlab.weizmann.ac.il/McPAS-TCR/, accessed March, 1^st^, 2022). The code to reproduce the results is available at https://github.com/SchubertLab/meTCRs. All reported models can be downloaded from https://doi.org/10.5281/zenodo.7113264.

## Notes

### Competing Interest Statement

The authors have declared no competing interest.

